# RNA evidence supports clinical stage antibody drug targets in oncology

**DOI:** 10.64898/2026.10.06.757024

**Authors:** Daniele Pozzobon, Nichal Khusial, Roberto Spreafico, Attila Csala

**Affiliations:** Genmab; Absci

## Abstract

**Background:** The selection of efficacious and safe antibody-based oncology targets depends on tumor specificity and normal-tissue expression, and remains challenging. Bulk RNA-seq can identify tumor-enriched surface genes, but may obscure tumor specificity and normal-cell expression patterns that are relevant to the therapeutic window. We evaluated whether single-cell RNA-seq-derived expression features improve retrospective prioritization of clinically pursued antibody targets.

**Methods:** We developed a bulk and single-cell RNA expression-based ranking framework across six solid tumor indications. 4,532 candidate surfaceome targets were scored using tumor *Specificity*, normal *Safety*, and blood *Sink* features derived from bulk RNA-seq, single-cell RNA-seq, or a combined modality. Ranking performance was evaluated against a curated validation set of tumor-associated antibody-based oncology targets using logistic regression, with odds ratios (ORs) reported per one-standard-deviation increase in score.

**Results:** Higher *Specificity* scores were associated with clinical-stage target selection across all modalities. In cross-indication meta-analysis, single-cell *Specificity* showed higher ORs than bulk *Specificity* across clinical phase thresholds, including Phase ≥ 1 (OR 2.56 versus 1.81), Phase ≥ 2 (OR 2.77 versus 1.88), Phase ≥ 3 (OR 3.53 versus 2.04) and *Approved* targets (OR 3.51 versus 1.74). Combined bulk-and-single-cell *Specificity* showed intermediate enrichment. *Specificity* score alone did not consistently show a significant single-cell advantage over bulk in paired indication-level comparisons, except for Phase ≥ 3 targets. After adding normal-cell *Safety* and blood *Sink* features, the single-cell general score showed the strongest enrichment across thresholds (Phase ≥ 1 OR 3.61, Phase ≥ 2 OR 4.16, Phase ≥ 3 OR 5.57, and Approved targets OR 4.25). This general score also showed consistently significant single-cell advantage over bulk in paired indication-level comparisons. *Safety* contributed most strongly in the single-cell modality, increasing meta-analytic ORs by an average of ∼36% across thresholds compared to *Specificity* alone. Performance varied by indication, with breast cancer showing weaker *Specificity*-based enrichment.

**Conclusion:** Expression-derived ranking features are retrospectively enriched among clinically pursued tumor-associated antibody targets. Single-cell evidence is most informative when tumor *Specificity* is combined with normal-cell *Safety* information, supporting its use as an interpretable component of early antibody target prioritization. These findings are in alignment with recent evidence that single-cell RNA-based cell-type specificity supports clinical-stage targets and demonstrate that expression-driven ranking can systematically recover clinically relevant oncology targets.

## Introduction

Clinical drug development remains to have high clinical failure rate to this day. Across therapeutic areas, only about 13% of candidates entering Phase 1 achieve approval, and in oncology the success rate is lower (Wong et al 2019). Analyses of clinical failures highlight that clinical failure is largely driven by insu>icient clinical e>icacy and unmanageable toxicity, alongside suboptimal drug-like-property, commercial and strategic factors (Hwang et al 2016, Sun et al 2022). For antibody-based therapeutics, one drug-related component of high clinical failure rates is the expression profile of the target. Insufficient tumor specificity, heterogeneous tumor expression, or expression in normal tissues can constrain the therapeutic window, thereby contributing to lack of efficacy or on-target toxicity. These constraints have motivated systematic, quantitative approaches to improve early-stage target prioritization (Fang et al 2024, Dann et al 2024, Li et al 2025). While human genetic evidence has shown a two-fold association with clinical success (Minikel et al 2024), many oncology programs lack interpretable genetic links, and oncology antibody targets are prioritized based on tumor-associated surface expression and therapeutic window rather than direct causal genetic evidence (Fang et al 2024, Dann et al 2024, Li et al 2025). Transcriptomic evidence therefore offers complementary, cell level evidence of target disease-relevance and safety.

Bulk RNA sequencing (bulk RNA-seq) expression profiles on tumor and healthy tissues are systematically collected in datasets such as The Cancer Genome Atlas (TCGA) and Genotype-Tissue Expression (GTEx), enabling tumor-normal expression comparisons across cancer types (The Cancer Genome Atlas Research Network 2013, GTEx Consortium 2020). Such bulk expression comparisons have been used to nominate tumor-associated surface targets in oncology, particularly for antibody drug conjugates (Fang et al 2024). More recent target prioritization frameworks have extended this concept by integrating additional evidence modalities, including single-cell RNA-seq (scRNA-seq) expression profiles (Deng et al 2024, Li et al 2025, Shraim et al 2025). Bulk profiles obscure cell-type heterogeneity and cannot distinguish tumor-cell expression from stromal, immune, or other normal-cell expression within a tissue, while scRNA-seq enables single-cell-level assessment of whether a target is enriched in tumor cells or shared with critical normal cell types (Liu et al 2024). However, to the best of our knowledge, no integrated framework has systematically compared how bulk and single-cell expression features relate to clinical-stage antibody target selection across multiple solid tumor indications.

For potential antibody oncology targets, expression-based prioritization commonly begins by identifying genes enriched in tumors relative to adjacent normal tissues. However, for antibody oncology targets, tumor specificity alone is unlikely to be sufficient. Targets must also have an acceptable normal-tissue expression profile, and expression in circulating blood compartments may reduce effective exposure or increased peripheral binding. We therefore investigated two related questions; first, whether cell-type-resolved single-cell expression-based evidence improves retrospective target prioritization compared with bulk expression-based evidence, and second, whether normal-tissue safety and blood sink features improve ranking beyond tumor specificity alone.

In this work, we introduce an expression-driven target ranking framework that integrates bulk and single-cell transcriptomic evidence to prioritize tumor-associated antibody drug targets with low normal tissue and blood expression across solid-cancer indications. The framework scores candidate genes along three interpretable features: tumor *Specificity*, normal Safety, and blood *Sink*. We compare bulk, single-cell, and combined single-cell-and-bulk evidence modalities, and evaluate whether highly ranked genes are enriched among clinically pursued tumor-associated antibody targets, providing a retrospective benchmark of how expression-driven target properties relate to oncology antibody target selection.

## Results

### Clinical antibody target landscape and ranking framework overview

We first asked whether expression-derived target features recover tumor-associated antibody targets that have been clinically pursued in solid tumors. Based on prior work showing that tumor expression and cell-type specificity are associated with clinical target selection (Dann et al, 2024, Shraim et al 2025, Zhang et al 2026), we hypothesized that scRNA-seq evidence would improve target prioritization over bulk RNA evidence by distinguishing tumor-associated targets from targets broadly expressed across healthy or non-malignant cell populations.

To test this hypothesis, we developed an expression-based target ranking framework across six solid tumor indications and compared three evidence modalities: bulk, single-cell, and a combined bulk-and-single-cell modality. Within this framework, each input candidate surfaceome gene was scored along three interpretable features: tumor *Specificity*, normal *Safety*, and blood *Sink*. *Specificity* captured enrichment in tumor relative to matched adjacent normal tissues, reflecting potential target tumor specificity. *Safety* captured low and restricted expression across healthy tissues or cell types, reflecting potential on-target toxicity. *Sink* captured expression in blood compartments, reflecting potential peripheral target binding or target-mediated drug disposition. These features were normalized and combined into a general ranking score, while *Specificity*-only rankings were used as the primary baseline for benchmarking. We evaluated the ranking performance against a validation set of curated clinically investigated antibody targets from oncology trials, restricted to tumor-specific or tumor-cell-associated targets. The validation set covered breast, colorectal, endometrial, head and neck, lung and ovarian cancers, and included targets across multiple antibody-based formats and clinical phases. This clinical landscape was heterogeneous, characterized by a diverse range of targets explored across multiple antibody-based formats and a low degree of success rate across development phases (**Fig. 1**). Notably, a relatively limited set of targets is repeatedly investigated across trials and formats, reflecting both biological relevance and historical prioritization strategies.

**Figure 1.**
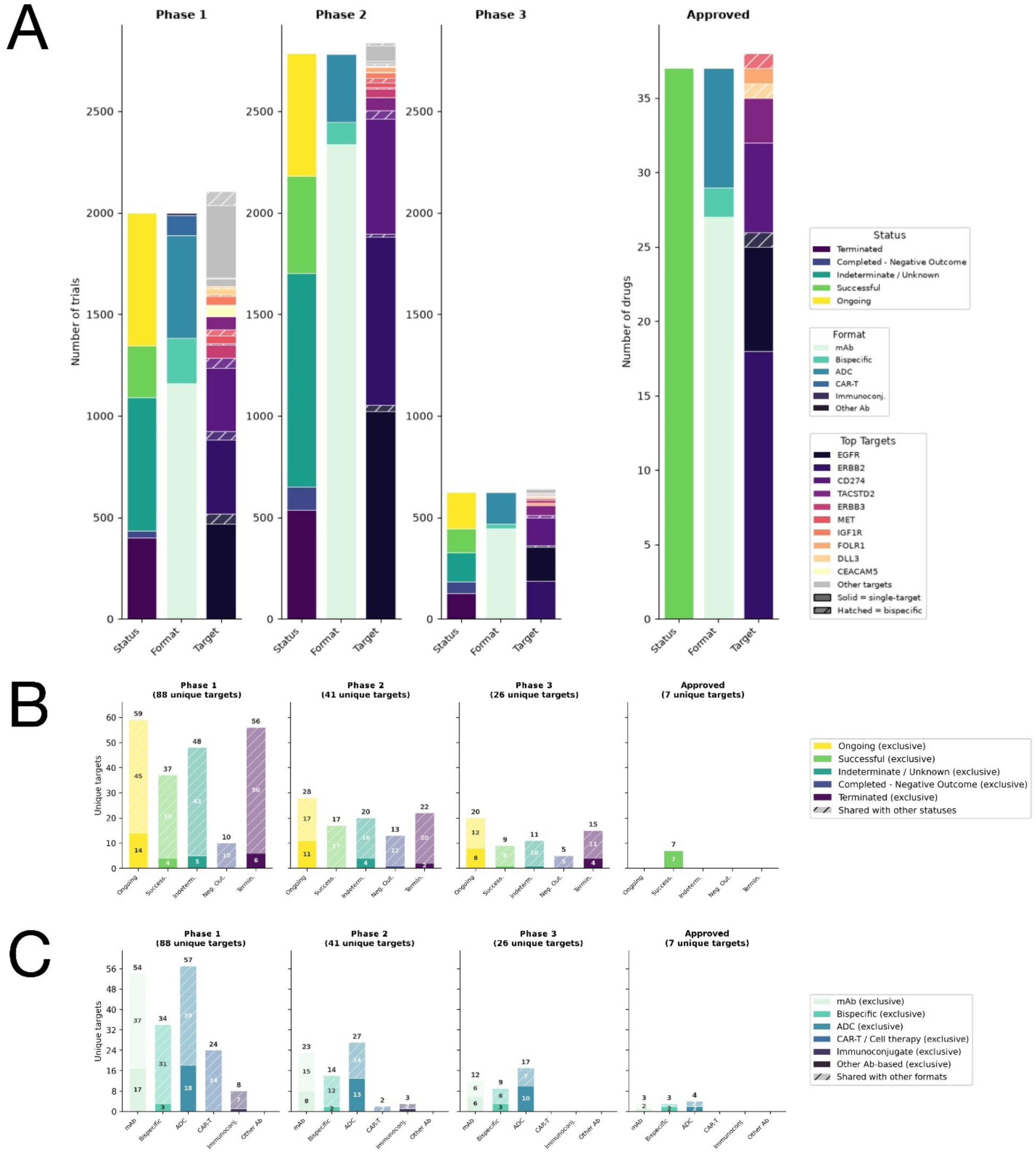
Landscape of antibody-based clinical trials and validated targets across development phases. **(A)** Stacked bar chart summarizing the number of trials per phase (1-3 and Approved) stratified by trial status, drug format, and top target gene; hatched segments denote bispecific constructs. **(B)** Unique targets by trial status per clinical phase targets (1-3 and Approved); solid bars represent targets exclusive to a single status, hatched bars represent targets shared across multiple statuses. **(C)** Unique targets by drug format per clinical phase targets (1-3 and Approved); solid and hatched bars follow the same exclusive/shared convention as in (B). Analysis restricted to surfaceome targets in antibody- and cell-based trials across six solid-tumor indications (ovarian, breast, lung, head and neck, colorectal, endometrial).

Given that tumor-specific expression has been previously identified as a primary determinant of target selection (Fang et al 2024, Dann et al 2024, Li et al 2025), we first tested whether the *Specificity* feature alone could capture a consistent signal underlying clinical target selection. We then extended this framework to evaluate the additional contribution of *Safety* and *Sink* features, assessing whether these features further improve target prioritization beyond *Specificity*. Across all indications, we performed a pooled logistic regression analysis to evaluate the association between the feature based ranking scores and clinical relevance. In total 201 clinically investigated targets (defined using Phase 1, 2, 3 or Approved targets) were retrieved from Citeline (Norstella Inc, New York, NY, USA; see **Methods**), with 106 removed as immune-associated targets (e.g. CD3-encoding genes, CD274, CD47 and HLA-G) or in general for not being tumor-associated, 10 targeted non-surfaceome proteins (e.g. secreted or soluble antigens), and 12 additional targets had all Phase I trials failed, which were thus removed. Hence, 73 targets were used as binary outcome labels, with standardized ranking scores over 4,532 surfaceome candidate target gene as the main predictor. Clinically relevant targets were evaluated separately at Phase ≥ 1, Phase ≥ 2, Phase ≥ 3 and Approved thresholds. The indication was included in the model as a covariate to account for baseline differences across tumor types. The Odds Ratio (OR) obtained from the logistic regression results represented the change in odds that a gene is a clinical-stage target for each one-standard-deviation increase in *Specificity*, *Safety* and *Sink* score, indicating clinical relevance of highly ranked targets.

To correct for mixed effects, we also performed a meta-analysis aggregation to obtain a pooled OR as the geometric mean across the six individual indications. Statistical significance for forest-plot ORs was reported using the two-sided Wald tests for the logistic regression score coefficient, unless otherwise stated. To assess the robustness of the results, we compared the observed 95% confidence interval (CI) and a permutation-derived null intervals generated by permuting validation labels. Paired comparisons between bulk and single-cell modalities were assessed using paired t-test on indication-level log(OR) estimates.

### Pan-indication association between expression Specificity and clinical target selection

Across all evidence modalities (bulk, single-cell, and combined bulk-and-single-cell), higher *Specificity* scores were associated with increased likelihood of clinical-stage target status (**Fig. 2A**). In the cross-indication meta-analytic analysis, bulk-derived *Specificity* showed significant enrichment across all phase thresholds (OR 1.81 for Phase ≥ 1, OR 1.88 for Phase ≥ 2, OR 2.04 for Phase ≥ 3 and OR 1.74 for Approved targets), and only Approved results showed overlap with the permutation based null-intervals, indicating in general strong and robust association between tumor-specific expression and clinical target selection. Single-cell-derived *Specificity* showed higher point estimates at each threshold (OR 2.56 for Phase ≥ 1, OR 2.77 for Phase ≥ 2, OR 3.53 for Phase ≥ 3 and OR 3.51 for Approved targets) and no overlap with null-intervals. Combined bulk-and-single-cell-based *Specificity* also showed positive enrichment, with effect sizes generally intermediate between the bulk-only and single-cell-only modalities.

**Figure 2.**
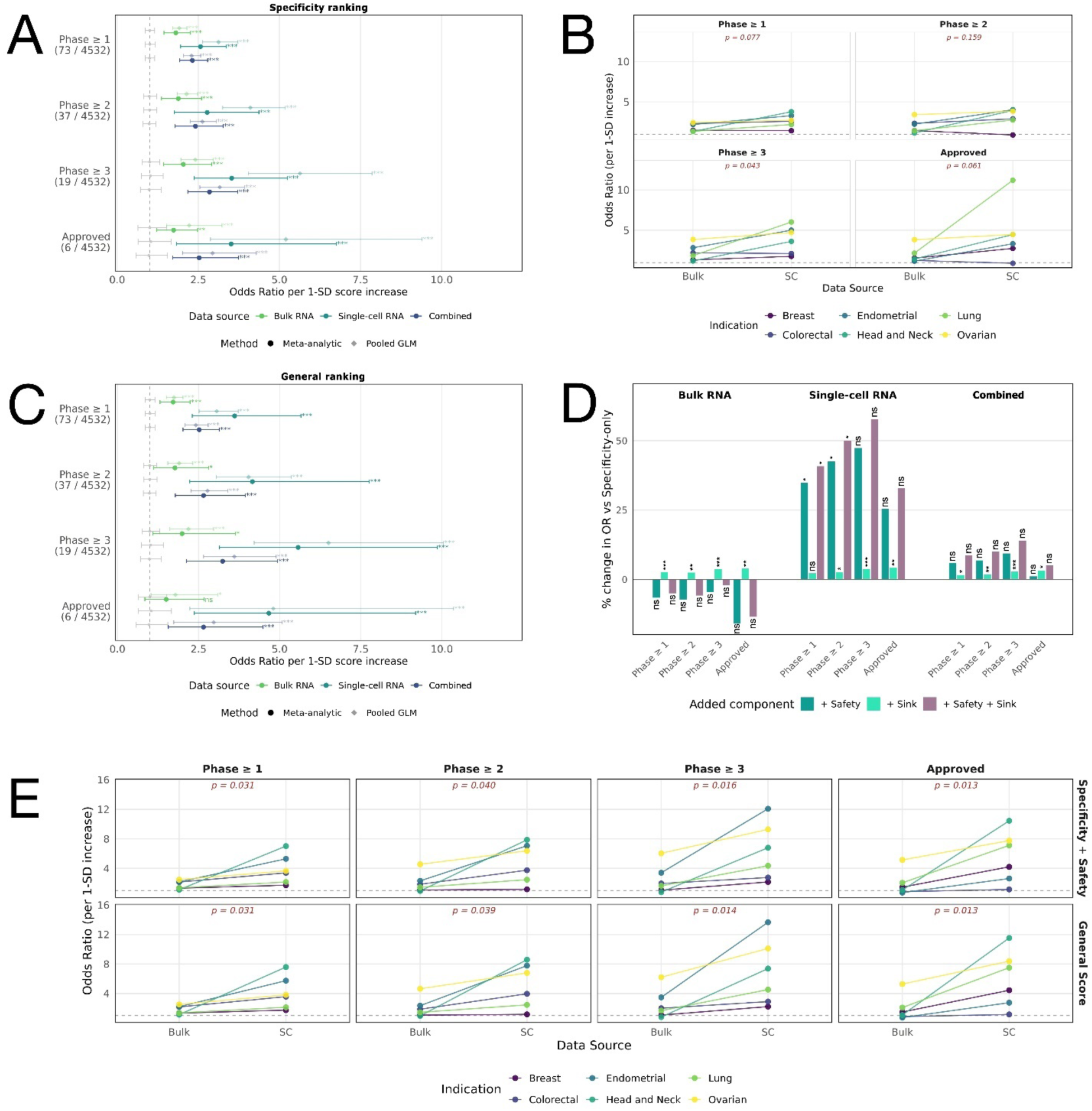
Expression-based ranking scores are associated with clinical-stage antibody target status across six solid indications. Odds ratios (ORs) quantify the association between ranking score and clinical-stage target status per one-standard-deviation increase in score, evaluated across 4,532 analyzed surfaceome genes in breast, colorectal, endometrial, head and neck, lung, and ovarian cancer. Where shown, solid circles represent cross-indication meta-analytic estimates calculated from indication-specific logistic regressions, and faded diamonds represent pooled logistic regression estimates with indication included as a covariate. Scores were derived from bulk RNA-seq (green), single-cell RNA-seq (teal), or combined bulk-and-single-cell evidence (dark blue). Clinical-stage labels were assigned from curated Citeline clinical trial data; targets were evaluated separately at Phase ≥ 1, Phase ≥ 2, Phase ≥ 3, and Approved thresholds. Asterisks denote nominal two-sided Wald-test significance for the logistic-regression score coefficient: * p < 0.05, ** p < 0.01, *** p < 0.001. Gray segments indicate 95% permutation-null intervals from 1,000 label shuffles and are shown as a robustness benchmark. **(A)** Forest plot of specificity-only score, stratified by phase threshold. Y-axis annotations show positive clinical-stage target labels out of analyzed gene set. **(B)** Paired comparison of per-indication ORs between bulk versus single-cell Specificity scores. Lines connect the same indication; red annotations show paired t-test p-values on log ORs. **(C)** As in (A), but for the general score combining Specificity, Safety, and Sink. **(D)** Percentage change in meta-analytic OR after adding Safety, Sink, or both features to the Specificity-only baseline. Significance shows paired t-tests on indication level log-ORs. **(E)** As in (B), but comparing bulk and single-cell ORs for Specificity + Safety and the general score.

The direction of e>ect was consistent across phase thresholds and modalities, with pooled logistic regression confirming significant associations in all cases. Confidence intervals were wider for the single-cell and combined modalities, reflecting greater variability across indications. But the overall pattern held even at the most stringent thresholds.

Permutation-derived null intervals confirmed robustness of the observed ORs, except for bulk-based modality for Approved targets.

We next compared bulk and single-cell *Specificity* scores directly across indications (**Fig. 2B**). Both modalities provided consistent enrichment, but the single cell advantage over bulk was heterogenous across indications. Single-cell *Specificity* had higher pan-indication point estimates at every threshold. However, paired indication-level comparisons reached statistical significance only for Phase ≥ 3 targets (paired t-test p-value 0.043), with non-significant trends for Phase ≥ 1 (p-value 0.077), Phase ≥ 2 (p-value 0.159), and Approved targets (p-value 0.061). Breast and colorectal cancers showed little or no improvement with the single-cell modality.

### Contribution of Safety and Sink features

We next evaluated whether adding normal *Safety* and blood *Sink* features improved ranking performance beyond tumor Specificity alone (**Fig. 2C-D**). In the meta-analytic analysis, the full general score, combining *Specificity*, *Safety* and *Sink*, remained significant across all phase thresholds and modalities. The strongest general-score enrichment was observed for the single-cell modality with ORs of 3.61 for Phases ≥ 1, 4.16 for Phases ≥ 2, 5.57 for Phases ≥ 3 and 4.25 for Approved targets. Bulk-based general scores showed smaller effects (ORs 1.72, 1.77, 2.00, and 1.94 respectively), while combined bulk-and-single-cell scores were intermediate (ORs 2.52, 2.65, 3.24, and 2.90, respectively). Permutation-derived null intervals were non-overlapping for 95% CIs of all results except for bulk-based ORs for Phase ≥ 2, Phase ≥ 3 and Approved targets, indicating robust results from single-cell based modalities.

The incremental contribution of *Safety* was most apparent in the single-cell modality (**Fig 2D**; **Suppl. Fig. S1**). Adding *Safety* to single-cell *Specificity* increased the meta-analytic OR by an average of 36% across phase thresholds, with the largest gain at Phase ≥ 3 (OR from 3.53 to 5.20, 47% increase). In contrast, adding *Safety* to bulk *Specificity* slightly decreased ORs (average 6% decrease) and combined scores showed intermediate gains (8% increase). Paired t-tests confirmed significance for single-cell *Safety* at Phase ≥ 1 and Phase ≥ 2, but not for bulk or combined Safety. *Sink* alone produced smaller and less significant OR changes than *Safety*. However, the full general score (combining Specificity, Safety and Sink) produced the largest single-cell gains (OR increases of 41-58% for Phases ≥ 1 and Phases ≥ 3, and 27% for Approved targets). Sink associated improvements reached statistical significance across all modalities except for Phase ≥ 1 targets in the single-cell case. These improvements were not indication specific, but consistent across almost all tumor types under investigation (as discussed in following case studies sections, also see **Suppl. Fig. S2-S4**), albeit with varying magnitude (**Suppl. Fig. S5**). Notably, larger increases in OR were observed in indications where *Specificity*-based rankings already showed strong enrichment (e.g. in head and neck cancer), suggesting that the *Safety* feature does not primarily compensate for weak *Specificity* but amplifies existing tumor-specific signal (**Suppl. Fig. S6**). It is worth noting that given the limited number of datapoints (six, one per each indication), paired t-tests were heavily dependent on patterns of consistency (rather than magnitude alone) (**Suppl. Fig. S5**), which returned non-significant p-values. The addition of more indications in the framework could result in stronger improvements. These results indicate that normal-cell *Safety* contributed more strongly than blood *Sink* to the improvement over *Specificity* alone, particularly when derived from single-cell data.

Consistent with this pattern, the relative advantage of single-cell over bulk evidence became clearer after incorporating *Safety* (**Fig. 2E**). In paired indication-level comparisons, single-cell *Specificity* alone did not consistently outperform bulk *Specificity* across Phases ≥ 1, Phases ≥ 2 and Approval thresholds (**Fig. 2B**). By contrast, *Specificity* + *Safety* scores showed higher single-cell than bulk ORs across all thresholds (paired t-test p-values: 0.031 in Phase ≥ 1, 0.040 in Phase ≥ 2, 0.016 in Phase ≥ 3 and 0.013 in Approved targets). A similar pattern was observed for the full general score (p-values of 0.031, 0.039, 0.014, and 0.013, respectively). Thus, the main single-cell advantage was not driven by tumor *Specificity* alone but emerged most clearly when tumor enrichment was combined with normal-cell *Safety* evidence.

### Ovarian Cancer Case Study

Given the indication-level heterogeneity observed in the pan-indication analyses, we next examined selected indications in more detail. In ovarian cancer, *Specificity* scores were consistently associated with clinical-stage target status across bulk, single-cell and combined bulk-and-single-cell modalities (**Fig. 3A**). For Phase ≥ 1 targets, all three modalities showed significant enrichment with ORs of 2.60 for bulk, 2.93 for single-cell, and 2.61 for combined *Specificity*. Similar yet modest effect sizes were observed for Phase ≥ 2 targets, with larger ORs increases for Phase ≥ 3 targets, especially for single-cell *Specificity* (OR 4.73). When the validation threshold was restricted to Approved targets, effect sizes were lower and uncertainty increased, consistent with the smaller number of positive labels (*FOLR1* and *ERBB2*). For Approved targets, bulk, single-cell and combined bulk-and-single-cell *Specificity* remained significant (OR 3.83, p-value 9.68e-04; OR 4.47, p-value 1.52e-03; OR 3.69, p-value 8.64e-4, respectively). Permutation-null intervals provided a consistent robustness benchmark for the Phase ≥ 1, Phase ≥ 2 and Phase ≥ 3 results, whereas the Approved target estimates showed greater overlap with the null-interval.

**Figure 3.**
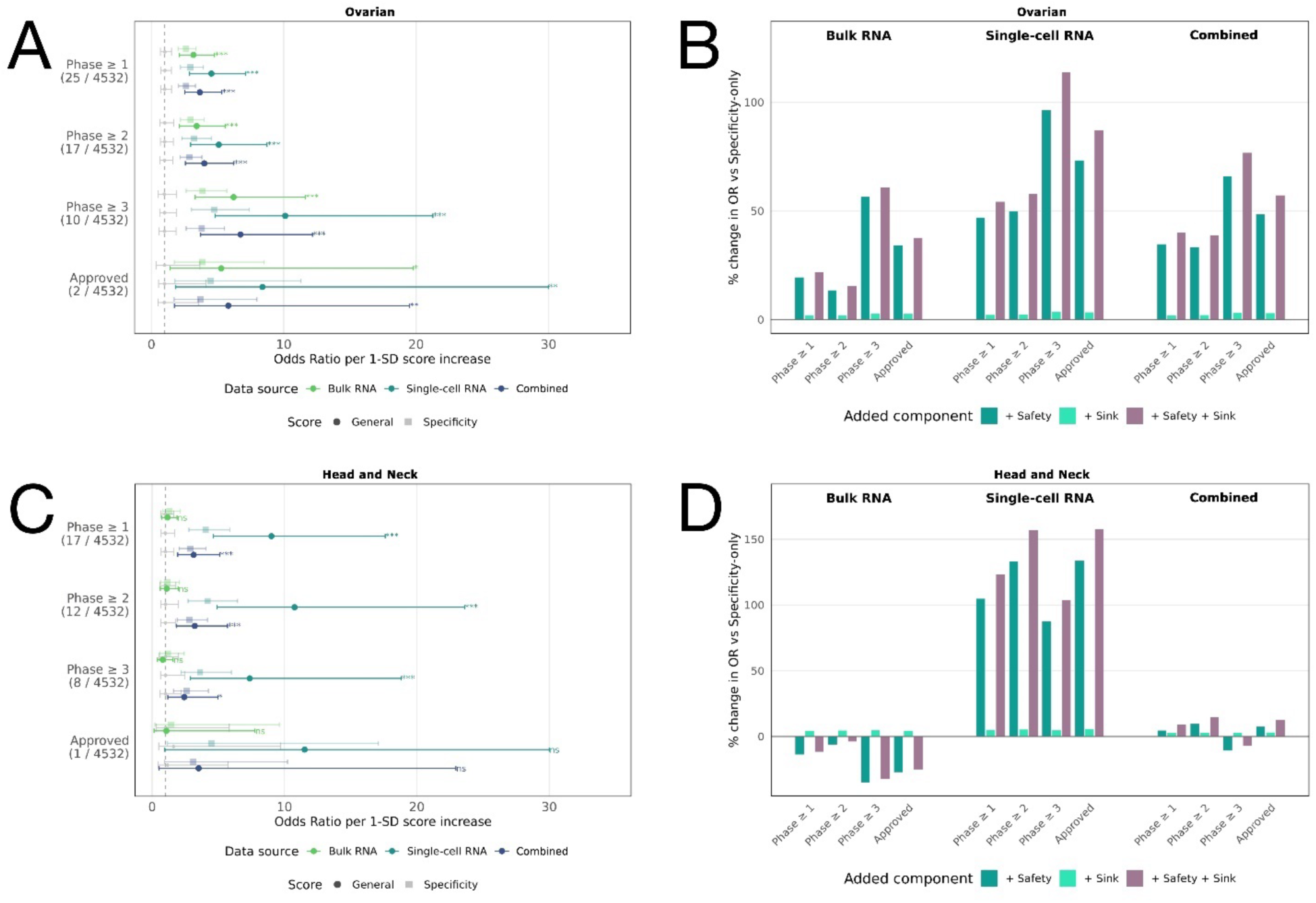
Indication-specific validation of expression-based ranking scores in ovarian and head and neck cancer. For each indication, odds ratios (OR) quantify the association between score and clinical-stage target status per one-standard-deviation increase in score, estimated using indication-specific logistic regression. Clinical-stage labels were assigned from curated Citeline clinical trial data filtered to the corresponding indication and evaluated separately at Phase ≥ 1, Phase ≥ 2, Phase ≥ 3, and Approved thresholds. Scores were derived from bulk RNA-seq (green), single-cell RNA-seq (teal), or combined bulk-and-single-cell evidence (dark blue). In forest plots, points indicate OR estimates and horizontal bars denote S5% Wald confidence intervals. Asterisks denote nominal two-sided Wald-test significance for the score coefficient: * p < 0.05, ** p < 0.01, *** p < 0.001. Gray segments indicate S5% permutation-null intervals from 1,000 label shuffles and are shown as a robustness benchmark. In bar charts, the zero line represents the Specificity-only baseline; each bar shows the percentage change in OR after adding Safety, Sink, or both features. Bar charts are descriptive and we refer the reader to the forest plots for inferential conclusions. **(A)** Forest plot of specificity score in ovarian cancer. **(B)** Percentage change in OR after adding Safety, Sink or both features in ovarian cancer. **(C)** Forest plot of specificity score in head and neck cancer. **(D)** Percentage change in OR after adding Safety, Sink or both features in head and neck cancer.

Adding *Safety* further improved the ovarian cancer ranking, particularly for the single-cell and combined bulk-and-single-cell modalities (**Fig. 3B**). Relative to the *Specificity*-only baseline, adding *Safety* increased single-cell ORs by 47.0%, 49.9%, 96.4%, and 73.3% for Phase ≥ 1, Phase ≥ 2, Phase ≥ 3, and Approved targets, respectively. The combined modality showed a similar pattern, with OR increases of 34.8%, 33.3%, 65.9%, and 48.4% across the same thresholds. In contrast, bulk *Safety* produced smaller and less consistent changes, with smaller gains at Phase ≥ 1 and Phase ≥ 2 (OR changes of +19.5% and +13.5% respectively), and larger increases for Phase ≥ 3 and Approved targets (OR change of +56.5% and +34.2%, respectively).

The general score, which added both *Safety* and *Sink* to *Specificity*, produced the largest OR increases in ovarian cancer. Compared to Specificity alone, the general score increased single-cell ORs by 54.3%, 57.8%, 113.8%, and 87.2% across Phase ≥ 1, Phase ≥ 2 and Phase ≥ 3 and Approved thresholds, respectively. Bulk and combined general scores also increased relative to their *Specificity*-only baselines, although the gains were smaller than for single-cell scores. Adding *Sink* alone produced modest but consistent OR increases approximately 2-5% across modalities and phase thresholds. Together, the ovarian cancer results support the pan-indication pattern: *Specificity* alone captured a strong clinical-stage target signal, while *Safety*, and a lesser extent *Sink*, improved enrichment most clearly when applied to single-cell-derived scores.

### Head and Neck Cancer Case Study

Head and neck cancer showed clear indication-specific advantages for single-cell derived ranking scores (**Fig. 3C-D**). For the *Specificity* score, single-cell and combined bulk-and-single-cell modalities were strongly associated with clinical-stage target status, whereas bulk *Specificity* did not show statistically significant enrichment. At the Phase ≥ 1 threshold, single-cell *Specificity* showed an OR of 4.04 (p-value < 0.001), and combined

*Specificity* showed an OR of 2.89 (p-value < 0.001), while bulk *Specificity* showed a smaller, non-significant association (OR 1.32, p-value 0.26). The same pattern was observed at the Phase ≥ 2, Phase ≥ 3 thresholds, with a single-cell *Specificity* OR of 4.19, combined *Specificity* OR of 2.82, and bulk *Specificity* OR of 1.15 for Phase ≥ 2 targets, and a single-cell *Specificity* OR of 3.62, combined *Specificity* OR of 2.62 and bulk *Specificity* OR of 1.20 for Phase ≥ 3 targets. For Approved targets, single-cell *Specificity* showed the strongest enrichment (OR 4.47, p-value 0.03), followed by the combined modality (OR 3.12, p-value 0.06), whereas bulk *Specificity* remained non-significant (OR 1.44, p-value 0.70).

Permutation-null intervals supported this pattern, with bulk estimates showing no separation from the null benchmark in any phase threshold, whereas the single-cell estimate showed separation from the null benchmark for Phase ≥ 1, Phase ≥ 2 and Phase ≥ 3 targets, suggesting robust results.

Adding *Safety* and *Sink* further strengthened the single-cell signal in head and neck cancer (**Fig. 3D**). Compared with the *Specificity*-only baseline, adding *Safety* increased single-cell ORs by 104.9% for Phase ≥ 1, 133.1% for Phase ≥ 2, 87.8% for Phase ≥ 3 and 133.8% for Approved targets. The full general score produced even larger single-cell gains, increasing ORs by 123.1%, 156.8%, 103.8% and 157.9% at the same thresholds, with general-score ORs reaching 9.02 for Phase ≥ 1, 10.76 for Phase ≥ 2, 7.38 for Phase ≥ 3, and 11.52 for Approved targets. In contrast, adding *Safety* to the bulk modality decreased ORs relatively to bulk *Specificity*, and the bulk general score remained non-significant across thresholds. For Phase ≥ 1, Phase ≥ 2 and Approved targets, combined bulk-and-single-cell scores improved modestly after adding *Safety* and *Sink*, but the gains were smaller than those observed for the single-cell-only modality (average OR change of 12.2%), while a decrease in OR was observed upon addition of combined *Safety* and *Sink* scores for Phase ≥ 3 targets (decrease of 7.24%). Together, these results suggest that, in head and neck cancer, the clinical-stage target signal was primarily captured by single-cell-based tumor *Specificity* and was further strengthened by single-cell-derived *Safety* evidence.

### Breast Cancer Case Study

Breast cancer represented an exception to the pan-indication pattern in which single-cell-derived rankings generally showed stronger enrichment than bulk-derived rankings. For the *Specificity*-only baseline, single-cell scores showed lower ORs (OR of 1.70) than bulk-and-single-cell combined scores (OR of 1.86) for Phase ≥ 1, and lower than the combined and bulk ORs in Phase ≥ 2 (single-cell Specificity OR of 1.34, combined *Specificity* OR of 1.58, and bulk *Specificity* OR of 1.56) (**Fig. 4A**). All three *Specificity* scores were significant for Phase ≥ 1, while single-cell *Specificity* was significant also for Approved targets (p-value 0.044). Permutation-null intervals showed limited separation from the null benchmark, consistent with a weaker *Specificity* signal in breast cancer than in the other case-study indications.

**Figure 4.**
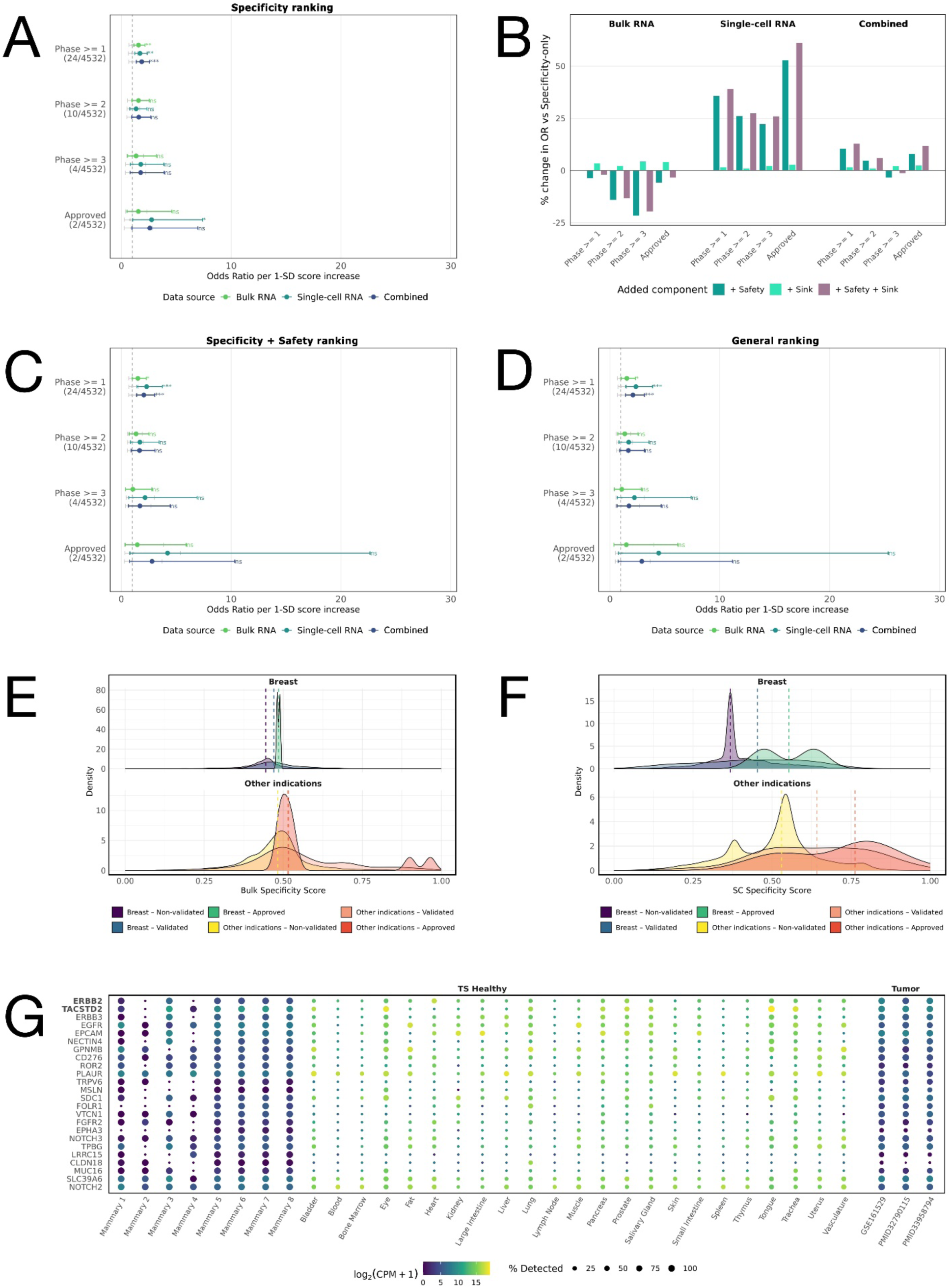
Breast cancer shows weaker specificity-based enrichment than other indications, with improvement after adding safety features. ORs were estimated by logistic regression per one-standard-deviation increase in score. Clinical-stage labels were assigned from curated Citeline clinical trial data and evaluated at Phase ≥ 1, at Phase ≥ 2, at Phase ≥ 3, and Approved thresholds. In forest plots, scores were derived from bulk RNA-seq (green), single-cell RNA-seq (teal), or combined bulk-and-single-cell evidence (dark blue). Points indicate OR estimates and horizontal bars denote S5% confidence intervals. Asterisks denote two-sided Wald-test significance for the score coeffect: * p < 0.05, ** p < 0.01, *** p < 0.001. Gray segments show S5% permutation-null intervals from 1,000 label shuffles. In density plots, targets are stratified into non-validated, validated clinical-stage targets (Phase 1–3), or Approved targets; dashed vertical lines indicate group medians. **(A)** Specificity-only forest plot for breast cancer. **(B)** Percentage change in OR after adding Safety, Sink, or both features to the Specificity-only baseline. Note that the bar chart is descriptive and we refer the reader to the forest plots for inferential conclusions. **(C)** Forest plot for the Specificity + Safety score. **(D)** Forest plot for the Specificity + Safety + Sink General score. **(E)** Bulk Specificity distributions for breast cancer compared with the pooled Specificity distributions in the other indications. **(F)** Single-cell Specificity distributions for breast cancer compared with the pooled Specificity Distributions in the other indications. **(G)** Single-cell pseudobulk expression of breast cancer (used for Specificity score) for clinical-stage targets across tumor and healthy reference datasets, showing expression level and detection frequency.

Adding *Safety* and *Sink* improved breast cancer ORs for the single-cell modality, but the absolute effect sizes remained modest (**Fig. 4B-D**). For single-cell scores, adding *Safety* increased ORs by 35.8%, 26.1%, 22.2%, and 52.9% across Phase ≥ 1, Phase ≥ 2, Phase ≥ 3, and Approved thresholds, respectively. However, single-cell *Specificity* + *Safety* retained significance only for Phase ≥ 1 targets (p-value 6.4e-03), together with bulk (p-value 0.46) and combined *Specificity* + *Safety* scores (p-value 3.2e-03). No significance was observed for all three modalities for Phase ≥ 2, Phase ≥ 3, and Approved targets. Upon further inclusion of the *Sink* scores, significant enrichment was shown again only for Phase ≥ 1 targets. *Sink* score alone returned only modest OR increases when added onto the *Specificity* baseline, with average OR change of +1.56%, +1.93% and +1.99% for bulk, single-cell and combined modalities, respectively. *Safety* and *Sink* scores together led to the largest enrichment only in single-cell ORs, with increases in OR of 39.4%, 27.5%, 26.0% and 61.1% for Phase ≥ 1, Phase ≥ 2, Phase ≥ 3 and Approved targets, respectively.

Across all modalities, Approved target general score ORs were larger (e.g. single-cell OR of 4.44) yet non-significant, indicating that the breast cancer Approved target set was not well captured by the expression-ranking framework.

To investigate this weaker enrichment, we compared *Specificity* score distributions for breast cancer clinical-stage targets and background genes with the corresponding distributions from the other indications. Breast cancer showed weaker separation between Phase ≥ 1 targets and background genes than the other indications, particularly for single-cell *Specificity*. In breast cancer, the effect size separation between clinically relevant Specificity scores and background score measured with the Cohen’s d statistic was 0.46 for bulk *Specificity* and 0.34 for single-cell *Specificity*, compared with 0.72 and 0.97, respectively, across all the remaining indications (**Fig. 4E-F**). Approved breast cancer targets showed higher *Specificity* scores than other clinical-stage targets, increasing the separation between background genes and Approved targets, with Cohen’s d of 0.53 for bulk and 1.52 for single-cell *Specificity*. The remaining indications presented similar Specificity scores between clinically-validated and Approved targets, reporting Cohen’s d statistics between non-validated and Approved targets of 1.05 and 1.50 for the bulk and single-cell modalities, respectively. The separation between Approved and background single-cell *Specificity* scores was found to be very similar in breast and in the remaining indications. However, single-cell specificity score for approved targets showed a clear bimodal distribution, suggesting different expression profiles and possibly different therapeutic window for *ERBB2* and *TACSTD2* (encoding for the approved targets HER2 and TROP2, respectively). However, the low number of Approved targets (n = 2) is likely to be a limiting factor when comparing score distributions between Approved vs. Non-validated genes.

Consistent with this pattern, many breast cancer clinical-stage targets showed comparable or higher expression in matched normal mammary gland samples than in tumor-cell pseudobulk profiles, including 18 out of 24 Phase ≥ 1 targets with negative single-cell log2FC values (e.g. *FOLR1* at –2.51, *MUC16* at –1.37 and *PLAUR* at –1.15) (**Fig. 4G**). Of the Approved targets, *TACSTD2* showed a strong negative Log2FC of -1.81, whereas *ERBB2* returned a weakly positive Log2FC of 0.39, in agreement with the bimodal single-cell *Specificity* distribution observed. Taken together, these results suggested that the lack of enrichment in the breast ranking outcomes was due to expression properties of the validation sets: in contrast to other indications, clinical-stage breast cancer targets did not exhibit a strong tumor-specific expression, which resulted in limited separation compared to background, non-clinical genes, leading to poor ranking performances. This has been further supported by the analysis of Specificity scores distributions, and the observation that many clinical-stage targets showed comparable or higher expression in normal mammary gland tissues. Moreover, these aspects suggested that, for breast cancer, clinical target selection may be less strongly aligned with broad tumor-versus-normal expression specificity than in other indications and may instead reflect factors not captured by this baseline expression framework, such as molecular subtype, target function, protein-level expression, drug format, or modality-specific therapeutic window.

## Methods

### Study design

We developed an expression driven framework to prioritize tumor-associated antibody drug targets with low normal tissue and blood expression. Analyses were performed both separately and pooled for six indications: breast, colorectal, endometrial, head and neck, lung and ovarian. For each indication, candidate target genes were ranked using three interpretable expression-derived features: tumor *Specificity* (enrichment in tumor tissue or tumor cells, depending on data modality), normal *Safety* (low or restricted expression across healthy tissues or cell types), and blood *Sink* (expression in circulating blood compartments). Ranking scores were computed using bulk RNA-seq and single-cell RNA-seq evidence modality and evaluated under three expression-evidence modalities: bulk, single-cell and combined bulk-and-single-cell. Ranking performance was benchmarked retrospectively against clinical-stage antibody targets, which were not used to train, tune or weight the ranking framework. Cross-indication performance was assessed using pooled logistic regression with indication as a covariate and by summarizing indication specific effects in a meta-analytic framework.

Throughout the Results, we use “modality” to refer to the expression evidence used for ranking: bulk, single-cell, or combined bulk-and-single-cell. We use “feature” to refer to the biological target property being scored for ranking: tumor *Specificity*, normal *Safety*, or blood *Sink*. A “ranking score” is the numeric score for a given modality-feature combination. A “ranking model” refers to the feature combination used to rank targets, such as *Specificity*-only, *Specificity* + *Safety*, or the general score combining *Specificity*, *Safety*, and *Sink*.

### Set of candidate targets

Because the intended therapeutic context is antibody-based targeting, the set of candidate targets was defined as genes encoding cell-surface or otherwise extracellularly accessible proteins (surfaceome). The surfaceome annotation was compiled from five public surfaceome or plasma membrane resources (Bausch-Fluck et al 2015, Bausch-Fluck et al 2018, da Cunha et al 2009, Fonseca et al 2016, Governa et al 2022) and one licensed annotation source from Euretos B.V (Leiden, the Netherlands). Genes were included if annotated as surface-associated in at least one source, resulting in 4,850 features. Gene identifiers were harmonized to HGNC symbols, deprecated symbols were updated where possible, and duplicates gene symbols were collapsed by summing gene counts. This produced 4,532 harmonized candidate surfaceome genes after indication-specific filtering for data availability.

### Bulk RNA-seq data processing

Bulk tumor expression data were obtained from The Cancer Genome Atlas (TCGA) for the six analyzed indications. Normal reference expression was obtained from Genotype-Tissue Expression (GTEx), with tissues selected to match or most closely represent the tissue of origin of each tumor type. TCGA and GTEx RNA-Seq by Expectation Maximization (RSEM) expected counts were retrieved from the UCSC Xena Toil Recompute Compendium, which provides uniformly processed TCGA and GTEx RNA-seq data and reduces technical bias between tumor and normal cohorts (Vivian et al 2017). Sample metadata were used to assign each sample to its tumor or normal tissue group.

For bulk tumor *Specificity* scoring, bulk tumor-normal differential expression analysis (DGEA) was used with TCGA and GTEX expression data. Expected counts were normalized using Trimmed Means of M-values (TMM) normalization in edgeR v3.64 (Chen et al 2025) and the tumor-normal differential expression was then estimated using the limma-voom v4.6 framework (Ritchie et al 2015). For each indication, the bulk tumor *Specificity* feature was the log2 fold-change (log2FC) comparing TCGA tumor samples with matched or relevant GTEx normal tissues.

For bulk normal tissue *Safety* scoring, bulk GTEx expression profiles were summarized across normal tissues at gene level based on Transcripts Per Million (TPM) normalized expression values. Median expression across tissues was used as a measure of normal expression, and the Tau index was used as a measure of tissue-restricted expression (Yanai et al 2005).

For bulk blood *Sink* scoring, gene expression in GTEx whole blood was summarized as mean expression per gene. Of note, whole blood bulk expression measurements are obtained directly from GTEx portal (https://gtexportal.org/home/), selecting the most recent release available (v11).

### Single-cell RNA-seq data processing

Single-cell RNA sequencing (scRNA-seq) tumor datasets were collected from publicly available resources. Where available, patient-matched adjacent normal samples were collected as additional reference controls. Healthy normal reference data were obtained from Tabula Sapiens v2 (Tabula Sapiens Consortium C Quake 2024). Cell-type annotations across tumor datasets were harmonized using MACE (Abdelaal et al 2025), with a reference dataset provided by Nebion AG (Zurich, Switzerland; now Immunai, New York). Malignant cells from primary tumor samples were selected as the tumor compartment for single-cell tumor *Specificity* modelling. A complete list of datasets used is provided in the Appendix. If any surfaceome genes were missing from some datasets, but the given gene was detected in at least one dataset, missing expression values were imputed by zero for the corresponding samples.

For single-cell *Specificity* scoring, we used log2FC obtained from DGEA applied to single-cell data as follows. Single-cell expression profiles of protein-coding genes were aggregated into pseudobulk samples, and lowly-expressed genes were filtered out using filterByExpr() with default parameters (limma v3.64, Ritchie et al 2015). Samples were used as the biological replicate unit when sample-level metadata were available, otherwise donor identifiers were used. Pseudobulk samples were normalized using edgeR’s TMM normalization and analyzed with limma-voom. When multiple samples from the same donor were present, donor-level correlation was modeled. Tumor-cell pseudobulk profiles were compared with patient-matched normal cells when available and with healthy reference cells otherwise. Of note, for head and neck cancer, single cell *Specificity* was estimated separately for anatomically matched regions, including larynx, oral cavity, oropharynx, and nasopharynx, because this disease group includes multiple normal tissues. Region-specific log2FC were then averaged to produce the final head and neck single-cell tumor *Specificity* estimate. Surfaceome genes filtered out because of low expression were imputed by 0 log2FC.

For normal expression *Safety* scoring, normal tissue expression profiles from Tabula Sapiens v2 were used using TPM-normalized expression values.

For blood *Sink*, the scores were computed based on expression in healthy blood cell populations (including granulocytes, platelets, monocytes, neutrophils, and erythrocytes) from Tabula Sapiens v2, using TPM-normalized expression values.

### Feature score definitions

All feature scores were designed so that larger values indicate more favorable target properties. Scores were min-max-normalized within each indication and data modality using:

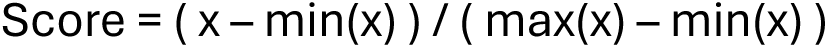

For each candidate gene, we computed three normalized features. Tumor *Specificity* was defined as tumor-enriched expression relative to relevant normal tissues. For the bulk modality, *Specificity* score was defined as the min-max-normalized bulk log2FC from TCGA versus GTEx differential expression. For the single-cell modality, *Specificity* score was defined as the min-max-normalized single-cell log2FC from tumor-cell pseudobulk profiles versus normal-cell pseudobulk profiles. For the combined bulk-and-single-cell modality, specificity score was defined by summing the normalized bulk and single-cell *Specificity* scores and min-max normalizing the sum.

Normal *Safety* score was defined as low expression in normal tissues combined with restricted expression across normal tissue or cell-type contexts. For the bulk modality, the *Safety* score was defined as the combined GTEx median expression across normal tissues with GTEx Tau index. For single-cell modality, the *Safety* score was defined as the combination of median expression across Tabula Sapiens cell-types with Tau index across tissue and cell-type groups. Specifically, median expression and Tau were each min-max-normalized, and the Safety score was computed as the min-max-normalized difference between normalized Tau and normalized median expression. Thus, genes with low normal expression and high tissue or cell-type restriction received higher *Safety* scores. The combined *Safety* score integrated the bulk and single-cell *Safety* components and was min-max-normalized.

Blood *Sink* score was defined as the expression in circulating blood compartments that may reduce effective target exposure or increase peripheral target binding. For the bulk modality, the *Sink* score was computed from GTEx whole blood mean expression. For the single-cell modality, *Sink* score was computed from Tabula Sapiens blood-associated populations, including granulocytes, platelets, monocytes, neutrophils, and erythrocytes. Expression values from blood compartments were min-max-normalized and multiplied by - 1 before final min-max normalization, so that lower blood expression produced a higher *Sink* score. The combined *Sink* score integrated bulk whole-blood expression and single-cell blood-cell expression and was min-max-normalized.

### General score aggregation

For each indication, ranking scores were computed under three evidence modalities: bulk, single-cell, and combined bulk-and-single-cell. Within each model, the general expression score was computed as a sum of tumor *Specificity*, normal *Safety* and blood *Sink* score, followed by min-max-normalization:

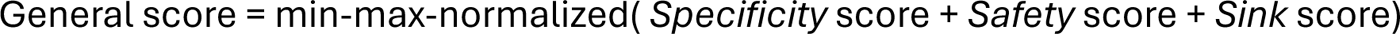

To evaluate the ranking results, we test whether highly ranked targets are enriched for clinical-stage oncology targets (i.e., approved and clinically investigated oncology targets targets).

Finally, to evaluate the contribution of individual feature groups, validation analyses were repeated for *Specificity*, *Specificity* plus *Safety*, and the full general score including *Specificity*, *Safety* and *Sink*. Genes were ranked in descending order, with higher scores indicating stronger predicted suitability as antibody targets.

### Clinical validation set

The clinical-stage antibody target validation set was obtained through curation of Trialtrove Citeline 2026 data (Norstella Inc, New York, NY, USA). All available oncology clinical trials were filtered for antibody-based drugs, including monoclonal antibodies, antibody drug conjugated, and bispecific or T-cell-engaging antibodies. Chimeric antigen receptor T-cell (CAR-T) therapies were also included, as the CAR antigen-recognition domain is derived from an antibody single-chain variable fragment and therefore targets the same class of accessible cell-surface antigens. Clinical trials with more than one experimental drug were excluded when the primary target could not be assigned unambiguously. For each retained trial, the primary molecular target or targets were mapped to HGNC gene symbols and linked to the relevant cancer indication.

Targets were classified whether the intended therapeutic antigen was primarily tumor-cell associated or instead reflected a non-tumor compartment, immune checkpoint. Since the ranking framework was designed to prioritize direct tumor-cell antigens, the validation set was restricted to tumor-specific or tumor-cell-associated targets. From 201 curated clinical-stage antibody targets, 95 were classified as tumor-specific, with 106 removed as immune-associated targets (e.g. CD3-encoding genes, CD274, CD47 and HLA-G) or in general for not being tumor-associated. Of the remaining targets, 10 were not annotated as encoding surface proteins, and 12 additional targets had all Phase I trials failed, which were thus removed. The 73 targets which were thus selected as validation set were labeled for clinical trial phases as follows:

• for each target, the maximum clinal phase reached in the relevant indication was recorded from Citeline;

• phase thresholds were analyzed separately as Phase ≥ 1, Phase ≥ 2, Phase ≥ 3 and Approved;

• a target was considered for a given phase threshold if it had either reached a higher phase than the threshold or its maximum phase equaled the threshold, with at least one ongoing or successful trial was recorded at that phase.

The 73 targets in the validation set were labeled for clinical trial phases as follows. For each target, the maximum clinal phase reached in the relevant indication was recorded from Citeline. Phase thresholds were analyzed separately as Phase ≥ 1, Phase ≥ 2, Phase ≥ 3 and Approved. A target was considered for a given phase threshold if it had either reached a higher phase than the threshold or its maximum phase equaled the threshold, with at least one ongoing or successful trial was recorded at that phase. Finally, the target labels were applied in an indication-specific manner, so that a target contributed a positive label only for the cancer indications in which it was clinically investigated.

### Statistical analysis

The ranking performance was evaluated using logistic regression against the clinical validation set. We used generalized linear models to fit the logistic regression models (stats v3.6.2, R core team 2026; logistf v1.24, Heinze et al. 2002). For each indication, ranking score, and phase threshold, clinical-stage target status was modeled as the binary outcome, with the corresponding standardized score as the predictor:

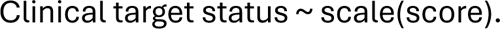

The odds ratio (OR), obtained from the logistic regression results, indicates how the odds of observing a clinical-stage target change as the ranking score increases. Because the ranking score is continuous, we first standardize it (mean = 0, standard deviation = 1), so that the resulting OR represents the change in odds that a gene is a clinical-stage target for each one-standard-deviation increase in the score.

We repeated this analysis using progressively stricter definitions of validation. Specifically, we defined clinical-stage targets as those that reached Phase 1, Phase 2, Phase 3 clinical trials (excluding targets that failed at the corresponding phase) or Approval. For each threshold, a separate logistic model was fitted and the OR together with its 95% confidence interval (CI) was reported. Results are summarized as forest plots showing the OR per one–standard-deviation increase in ranking score for each model variant (bulk, single-cell, and the combined bulk-and-single-cell modalities). Separate models were fitted for bulk, single-cell, and combined bulk-and-single-cell scores, and for Phase ≥ 1, Phase ≥ 2, and Phase ≥ 3 and Approved labels. In the case of very few validated targets (e.g., a single approved target), quasi-complete separation can prevent maximum likelihood estimation from converging. In such cases, Firth’s penalized likelihood was used (Firth, 1993, Heinze et al. 2002).

To assess pan-indication performance, we used two complementary approaches. First, we fitted logistic regression models with clinical-stage status as the outcome and the corresponding standardized *Specificity* score as predictor, which refer to as pooled logistic regression results. In this model we included indication as a covariate to account for baseline differences across tumor types:

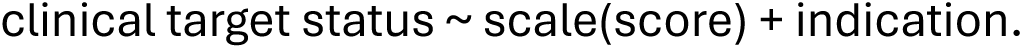

Second, we computed cross-indication summary effects from the indication-specific logistic regression results, which we refer to as the meta-analytic results. For each score type and phase threshold, per-indication log ORs were averaged across indications, and 95% CIs were calculated from the standard error of the mean log ORs across indications. These cross-indication meta-analytic summaries were used to assess whether enrichment was consistent across tumor types.

Unless otherwise stated, statistical significance for logistic-regression ORs was assessed using two-sided Wald test of the score coefficient. The asterisks shown in forest plots showed nominal Wald-test significance. In all forest plots, large 95% CI upper bounds have been clipped to an OR of 30 for visualization purposes.

Additionally, we further assessed the robustness of these results by generating OR null-distributions using a permutation-based approach. Within each indication, the null-distributions were obtained by randomly permuting the clinical association labels from the validation set and refitting the logistic regression models to the permuted data (n = 1,000 iterations). For these null-distributions, empirical 95% null intervals and higher quantiles were computed. Permutation-null intervals were used as a robustness benchmark to evaluate whether observed OR estimates were separated from label-randomized expectations.

To assess the contribution of individual feature groups (*Specificity*, *Safety*, and *Sink*), validation analyses were repeated for *Specificity* only, *Specificity* plus *Safety*, *Specificity* plus *Sink* and the full general score combining *Specificity*, *Safety*, and *Sink*. Feature contributions and modality differences were assessed using paired t-tests on indication-level log ORs. These paired tests were used only to evaluate whether ORs changed consistently across indications after adding a feature or comparing modalities. Because several indication-phase combinations contained few Phase 3 and Approved targets, sparse-data estimates, especially for *Sink*, were interpreted cautiously.

All validation analyses were retrospective and exploratory. Clinical-stage status was used only for benchmarking and was not used to train, tune, or weight the ranking model.

## Discussion

In this study, we investigated two related questions: first, whether cell-type-resolved single-cell expression evidence improves retrospective target prioritization compared with bulk expression-based evidence, and second, whether normal *Safety* and blood *Sink* features improve ranking results beyond tumor specificity alone.

To address the first question, we generated matched bulk-only, single-cell-only, and combined bulk-and-single-cell scores across *Specificity*, *Safety* and *Sink* feature combinations and evaluated their association with clinical-stage tumor-associated antibody target status using logistic regression and cross-indication meta-analytic aggregation. For *Specificity* alone, single-cell derived scores showed higher OR point estimates compared to bulk-derived scores across phase thresholds, although the paired indication-level comparison showed significance only for Phase ≥ 3 targets. However, when normal tissue derived *Safety* feature was added, single-cell derived ranking showed significantly higher enrichment than bulk-derived rankings across all phase thresholds, and the same pattern was observed for the full General score combining *Specificity*, *Safety* and *Sink*. These results suggest that cell-type-resolved single-cell expression evidence can improve retrospective target prioritization compared with bulk-expression derived evidence, particularly when the framework incorporates normal *Safety* features. However, the magnitude of improvement varied by indication, and breast cancer remained an important exception in which single-cell Specificity alone did not improve enrichment.

Furthermore, we have found no enrichment for the bulk-based scores in head and neck cancers, with ORs consistently around 1 both for the *Specificity*-only and General score rankings. This could be explained by the lack of matched healthy tissues from GTEx for DGEA, with only “salivary gland” available as most relevant tissue for bulk-based Specificity scoring. Alternatively, the anatomical and cellular heterogeneity of head and neck cancer may make cell-type-resolved expression particularly informative for tumor-associated target identification in this indication. Indeed, our single-cell Specificity analysis accounted for this heterogeneity by performing separate tumor-normal comparisons across anatomically matched regions. At present, the availability of anatomically matched bulk normal-tissue expression data for head and neck cancer is limited. As more appropriate reference datasets become available, future analyses will be needed to determine whether improved tissue matching strengthens the performance of bulk-based rankings in this indication.

We then asked whether adding normal tissue *Safety* and blood *Sink* features improved ranking beyond tumor *Specificity* alone. Based on the meta-analytic analysis results, the clearest evidence for increased ranking performance came from *Safety*, particularly in the single-cell modality. Adding *Safety* to single-cell *Specificity* increased the meta-analytic OR across all phase thresholds, with gains of approximately 35% for Phase ≥ 1, 43% for Phase ≥ 2, 47% for Phase ≥ 3, and 20% for Approved targets. These increases were supported by paired indication-level comparisons, indicating that the improvement was not driven only by a single tumor type (with large sample size). The full General score, which combined *Specificity*, *Safety* and *Sink*, showed a similar trend with larger improvement for single-cell-based ranking, with OR gains of approximately 27-58% across thresholds (also highest for Phase ≥ 3 targets).

In contrast, *Sink* alone contributed smaller but incremental gains. Across modalities and phase thresholds, adding *Sink* to *Specificity* generally produced only modest changes in OR. Although some *Sink* comparisons reached nominal significance, the effect sizes were small than those observed for *Safety*. This suggested that blood associated expression may add some information to ranking, but its contribution is secondary relative to broader normal-cell *Safety* information.

The bulk and combined modalities showed more modest and variable gains. For bulk-based scores, *Safety* and *Sink* produced negative or small changes in OR, respectively, and the full general score did not improve ranking. The combined bulk-and-single-cell model showed intermediate behavior, where *Safety* and the General score improved ORs relative to *Specificity* alone, but the gain was smaller than single-cell only results. Together, these results suggest that the added value of normal-expression features is strongest when derived from cell-type-resolved single-cell data, where normal-cell *Safety* evidence can refine tumor *Specificity* by penalizing targets with broader healthy-cell expression.

These findings support the conclusion that ranking based on tumor *Specificity* alone can be improved for antibody target prioritization: adding *Safety* features improves retrospective enrichment for clinically pursued targets, especially for models with single-cell evidence.

However, the evidence for *Sink* is weaker and should be interpreted as exploratory.

Of note, we have used processed clinical data to obtain our validation set for enrichment tests. The tested expression-based ranking models rely on the assumptions that antibody drug targets have inherent properties modelled by expression data that provides evidence for drug development success and failures. At the same time, clinical target selection is influenced by a range of factors beyond intrinsic target properties, including drug format, mechanism of action, development feasibility, commercial and strategic factors which are beyond the scope of our expression-driven framework. Future work is needed to expand the current target ranking framework to include additional drug development relevant aspects for target prioritization. For instance, additional components could be designed to account for different antibody formats, yielding novel target identification and prioritization outcomes for each drug format or mechanism of action of interest.

Our findings are broadly consistent with emerging evidence that expression-derived and cell-type-resolved features can inform therapeutic target prioritization. Dann et al. (2024) evaluated two complementary features: cell type specificity in healthy disease-relevant tissue, conceptually related to our normal *Safety* score, and disease cell specificity, which parallels our tumor *Specificity*. They found that both features were enriched in targets entering clinical development, and that cell type specificity was robustly associated with early-stage progression (Phase 1-2), whereas associations with late-stage development and drug approval were not significant. In our oncology-focused analysis, expression-based enrichment extended across all clinical-stage thresholds, including Approved targets. More recently, Zhang et al. (2026) derived cell-type specificity and expression bimodality features from healthy-tissue single-cell atlases. They reported significant associations with multiple clinical outcomes, including phase progression, trial endpoint success and reduced adverse event rates, extending to late-stage development, consistent with our finding but in contrast to the early-stage pattern observed by Dann et al. notably, the healthy-tissue features evaluated by Zhang et al. are conceptually more closely related to our normal *Safety* features than to our tumor *Specificity*. Shraim et al. (2025) similarly demonstrated the utility of integrating tumor expression with normal-tissue and other target features for immuno-oncology prioritization. Our framework complements these approaches by directly comparing matched bulk and single-cell evidence across the same features and indications, and by quantifying the incremental contribution of normal *Safety* features beyond tumor *Specificity* alone. However, differences in target scope, expression data, scoring strategies, and validation endpoints across these studies preclude direct quantitative comparison. Together, these studies support the value of expression-based evidence for target prioritization, while our results further suggest that cell-type-resolved normal-expression information and tumor tissue- and cell-type specific expression provides complementary signals for antibody target identification and prioritization.

Collectively, our work suggested that single-cell RNA-seq data improved target prioritization compared to bulk RNA-seq alone. In addition, incorporating safety-related features substantially increased enrichment for clinical phase targets, highlighting the importance of considering o>-tumor expression across healthy tissues and cell types during target selection. In contrast, sink-related features led to more modest improvements, suggesting that blood-associated target expression may play a smaller role in identifying clinically relevant targets than broader safety-associated expression patterns.

## Conclusion

Across six solid indications, tumor cell enriched RNA expression, especially single cell RNA evidence, is retrospectively enriched among clinically tested tumor-associated antibody targets. Expression *Specificity* is a generally robust retrospective signal for clinical-stage tumor-associated antibody target selection. Single-cell evidence often strengthens this signal by resolving tumor-cell expression, but performance is heterogeneous. Adding normal expression *Safety* features can improve enrichment, particularly for single-cell modality, while blood *Sink* contributes smaller and less consistent gains. Together, the framework provides an interpretable expression-based baseline for early antibody target prioritization in oncology drug discovery, and a foundation for incorporating additional target properties beyond expression.

## Supplementary Material

**Supplementary Figure S1.**
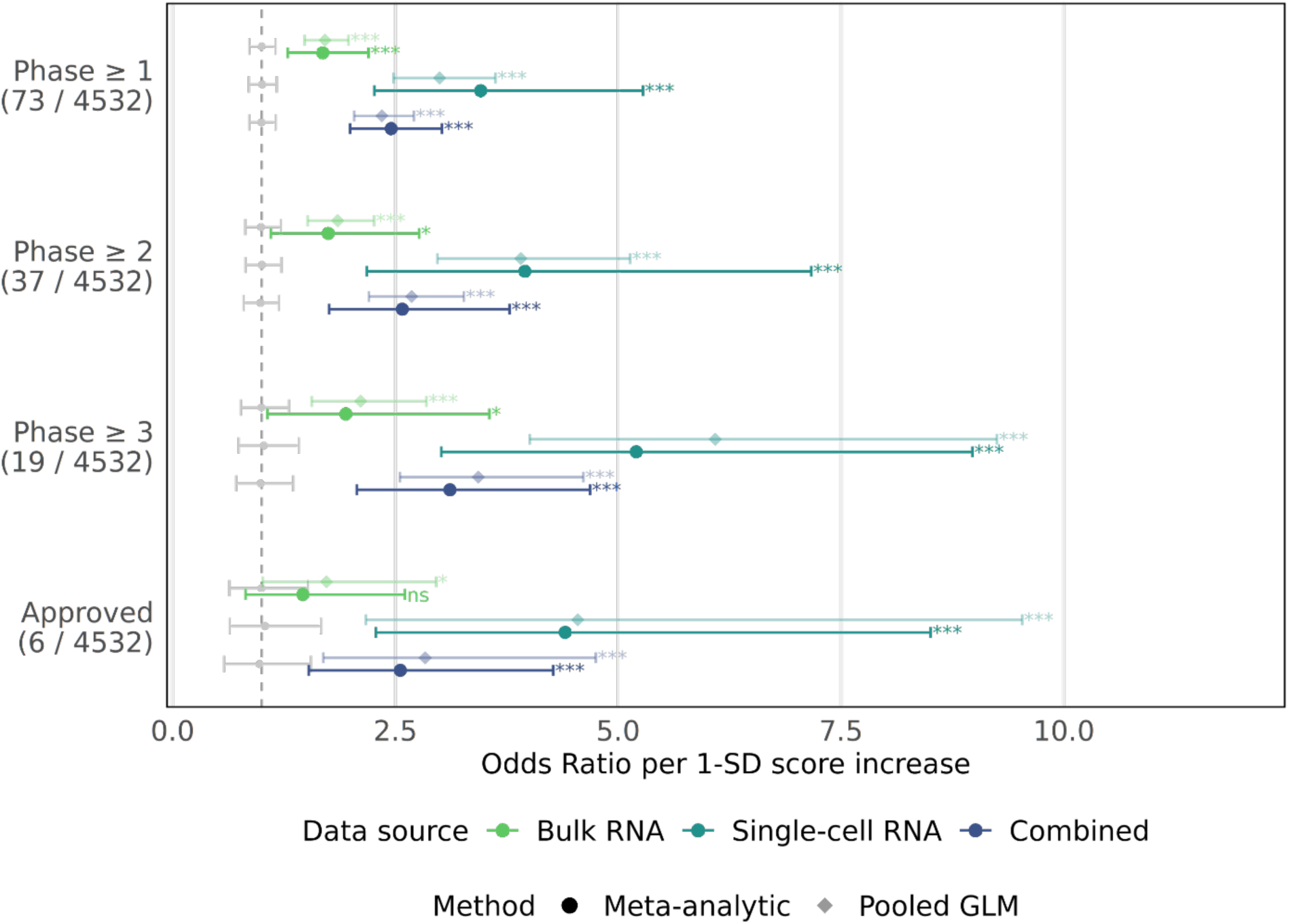
Association of Specificity + Safety ranking scores with clinical-stage antibody target status across six solid tumor indications. Odds ratios (ORs) quantify the association between the Specificity + Safety ranking score and clinical-stage target status per one-standard-deviation increase in score across breast, colorectal, endometrial, head and neck, lung, and ovarian cancer. Solid circles represent cross-indication meta-analytic estimates calculated from indication-specific logistic regressions, and faded diamonds represent pooled logistic regression estimates with indication included as a covariate. Scores were derived from bulk RNA-seq (green), single-cell RNA-seq (teal), or combined bulk-and-single-cell evidence (dark blue) and evaluated separately at Phase ≥ 1, Phase ≥ 2, Phase ≥ 3, and Approved thresholds. Y-axis annotations indicate the number of positive clinical-stage target labels out of 4,532 analyzed surfaceome genes. Asterisks denote nominal two-sided Wald-test significance for the logistic-regression score coefficient: * p < 0.05, ** p < 0.01, *** p < 0.001. Gray segments indicate 95% permutation-null intervals from 1,000 label shuffles.

**Supplementary Figure S2.**
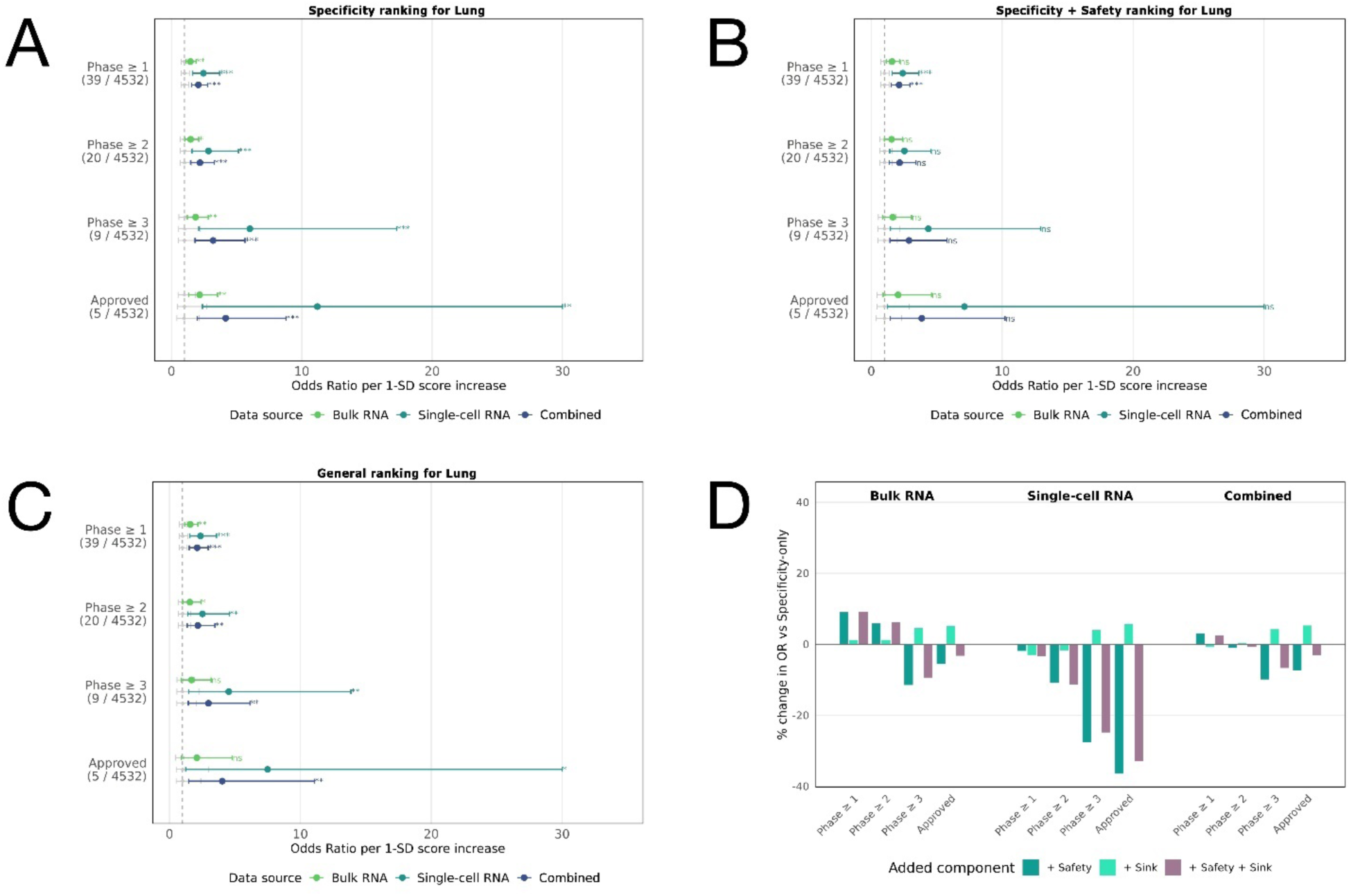
Indication-specific validation of expression-based ranking scores in lung cancer. ORs were estimated by logistic regression per one-standard-deviation increase in score. Clinical-stage labels were assigned from curated Citeline clinical trial data and evaluated at Phase ≥ 1, at Phase ≥ 2, at Phase ≥ 3, and Approved thresholds. In forest plots, scores were derived from bulk RNA-seq (green), single-cell RNA-seq (teal), or combined bulk-and-single-cell evidence (dark blue). Points indicate OR estimates and horizontal bars denote 95% confidence intervals. Asterisks denote two-sided Wald-test significance for the score coeffect: * p < 0.05, ** p < 0.01, *** p < 0.001. Gray segments show 95% permutation-null intervals from 1,000 label shuffles. **(A)** Specificity-only forest plot for lung cancer. **(B)** Forest plot for the Specificity + Safety score. **(C)** Forest plot for the Specificity + Safety + Sink General score. **(D)** Percentage change in OR after adding Safety, Sink, or both features to the Specificity-only baseline. Note that the bar chart is descriptive and we refer the reader to the forest plots for inferential conclusions.

**Supplementary Figure S3.**
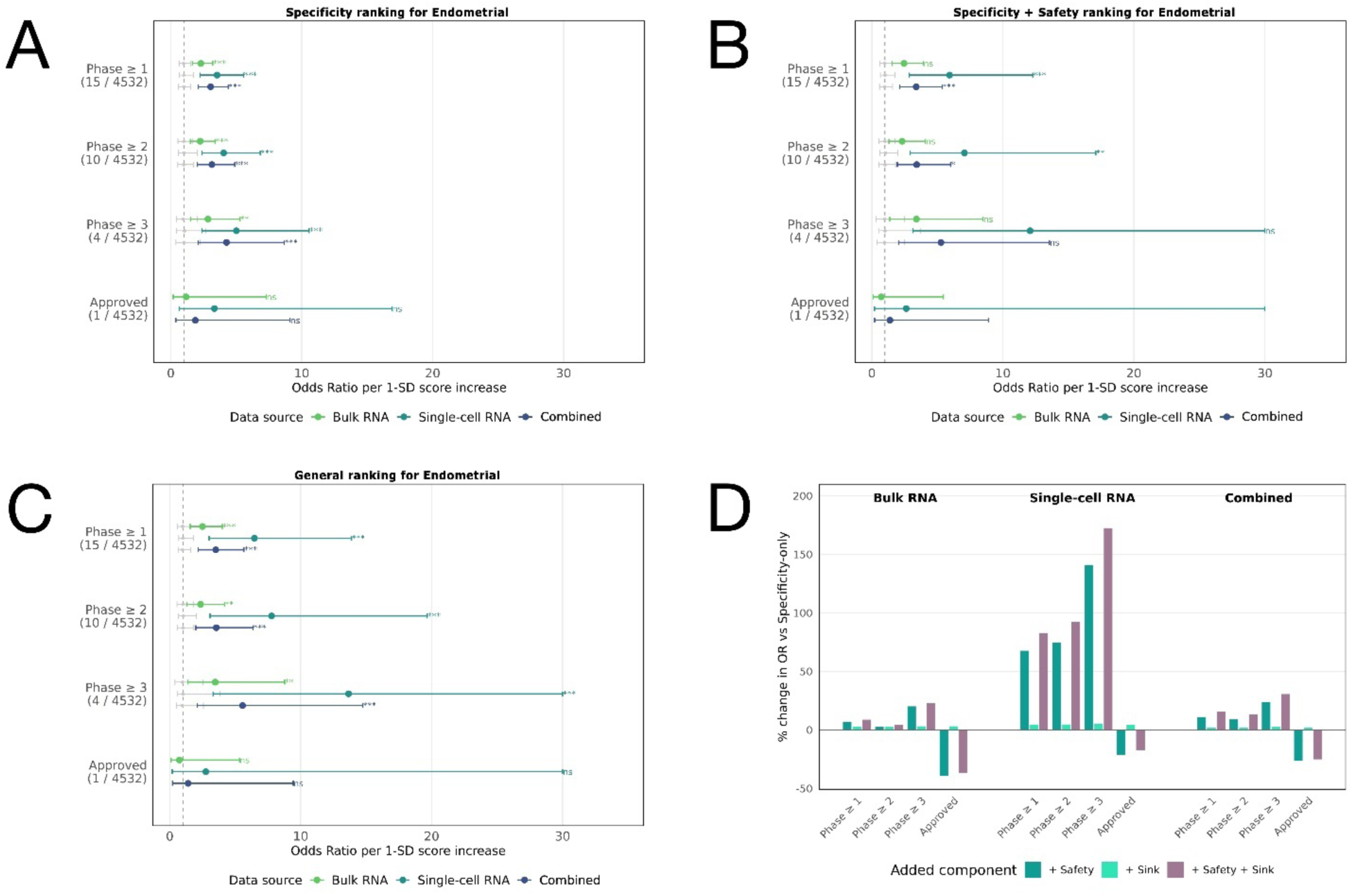
Indication-specific validation of expression-based ranking scores in endometrial cancer. ORs were estimated by logistic regression per one-standard-deviation increase in score. Clinical-stage labels were assigned from curated Citeline clinical trial data and evaluated at Phase ≥ 1, at Phase ≥ 2, at Phase ≥ 3, and Approved thresholds. In forest plots, scores were derived from bulk RNA-seq (green), single-cell RNA-seq (teal), or combined bulk-and-single-cell evidence (dark blue). Points indicate OR estimates and horizontal bars denote 95% confidence intervals. Asterisks denote two-sided Wald-test significance for the score coeffect: * p < 0.05, ** p < 0.01, *** p < 0.001. Gray segments show 95% permutation-null intervals from 1,000 label shuffles. **(A)** Specificity-only forest plot for endometrial cancer. **(B)** Forest plot for the Specificity + Safety score. **(C)** Forest plot for the Specificity + Safety + Sink General score. **(D)** Percentage change in OR after adding Safety, Sink, or both features to the Specificity-only baseline. Note that the bar chart is descriptive and we refer the reader to the forest plots for inferential conclusions.

**Supplementary Figure S4.**
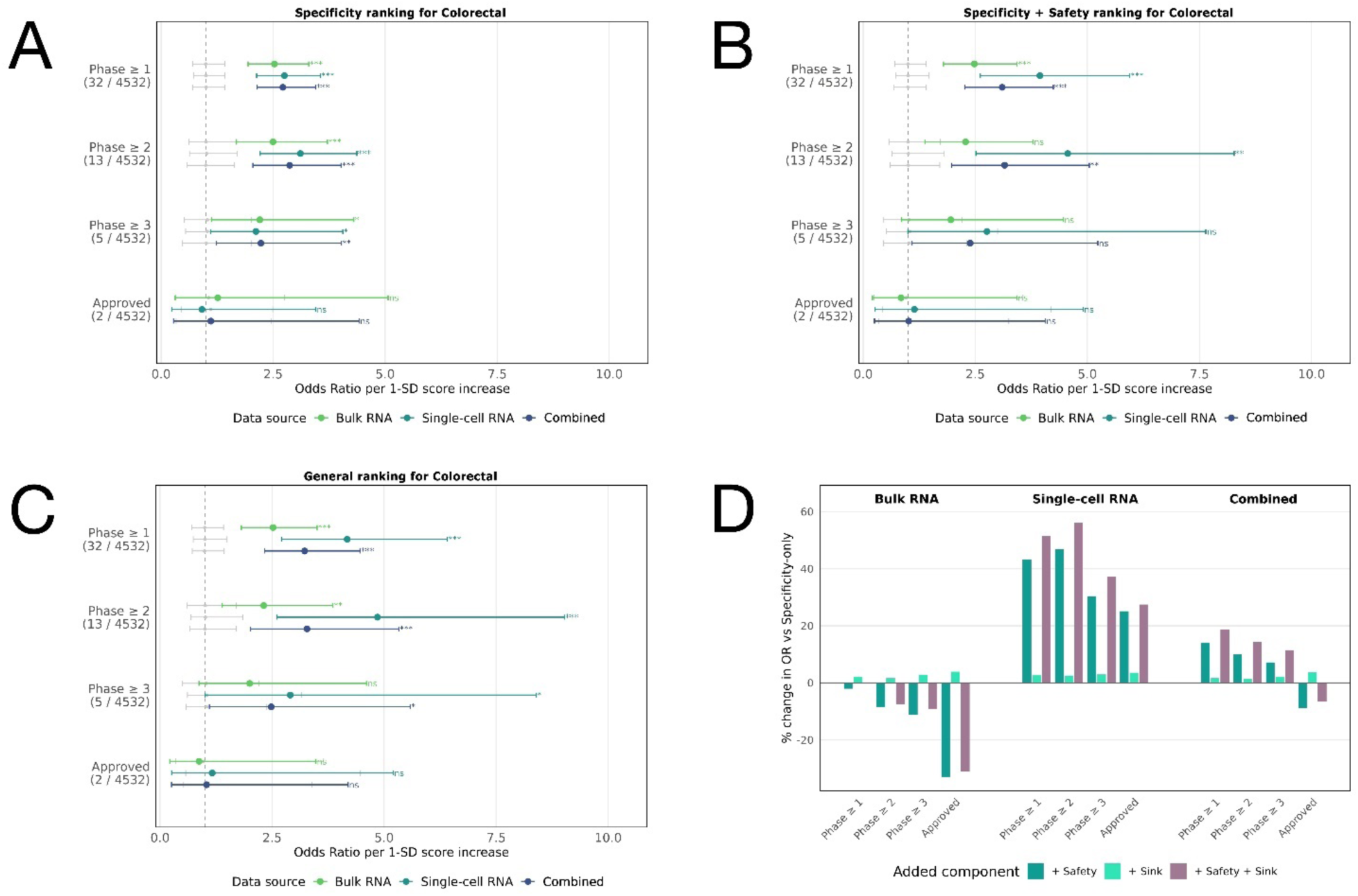
Indication-specific validation of expression-based ranking scores in colorectal cancer. ORs were estimated by logistic regression per one-standard-deviation increase in score. Clinical-stage labels were assigned from curated Citeline clinical trial data and evaluated at Phase ≥ 1, at Phase ≥ 2, at Phase ≥ 3, and Approved thresholds. In forest plots, scores were derived from bulk RNA-seq (green), single-cell RNA-seq (teal), or combined bulk-and-single-cell evidence (dark blue). Points indicate OR estimates and horizontal bars denote 95% confidence intervals. Asterisks denote two-sided Wald-test significance for the score coeffect: * p < 0.05, ** p < 0.01, *** p < 0.001. Gray segments show 95% permutation-null intervals from 1,000 label shuffles. **(A)** Specificity-only forest plot for breast cancer. **(B)** Forest plot for the Specificity + Safety score. **(C)** Forest plot for the Specificity + Safety + Sink General score. **(D)** Percentage change in OR after adding Safety, Sink, or both features to the Specificity-only baseline. Note that the bar chart is descriptive and we refer the reader to the forest plots for inferential conclusions.

**Supplementary Figure S5.**
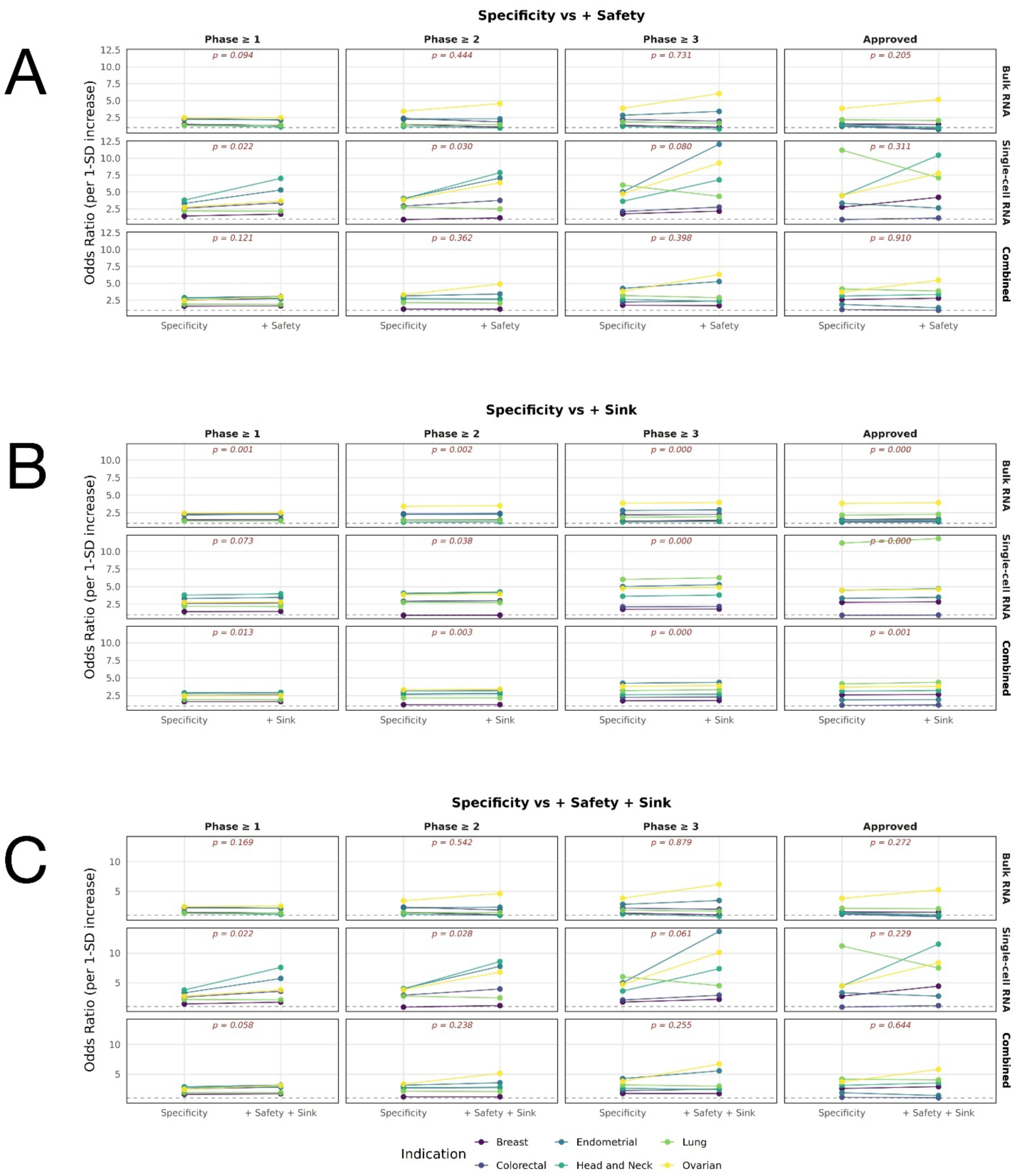
Indication-level changes in ranking enrichment after addition of Safety and Sink features to the Specificity-only baseline. Paired plots show indication-specific odds ratios (ORs) for clinical-stage antibody target status before and after addition of individual ranking features, evaluated separately for bulk RNA-seq, single-cell RNA-seq, and combined bulk-and-single-cell evidence. ORs quantify the association between ranking score and clinical-stage target status per one-standard-deviation increase in score and are shown separately for Phase ≥ 1, Phase ≥ 2, Phase ≥ 3, and Approved target thresholds. Lines connect OR estimates for the same indication between ranking models; colors denote breast, colorectal, endometrial, head and neck, lung, and ovarian cancer. Red annotations show paired t-test p-values calculated from indication-level log(OR)s. The horizontal dashed line indicates OR = 1. **(A)** Comparison of Specificity-only and Specificity + Safety rankings. **(B)** Comparison of Specificity-only and Specificity + Sink rankings. **(C)** Comparison of Specificity-only and General rankings combining Specificity, Safety, and Sink.

**Supplementary Figure S6.**
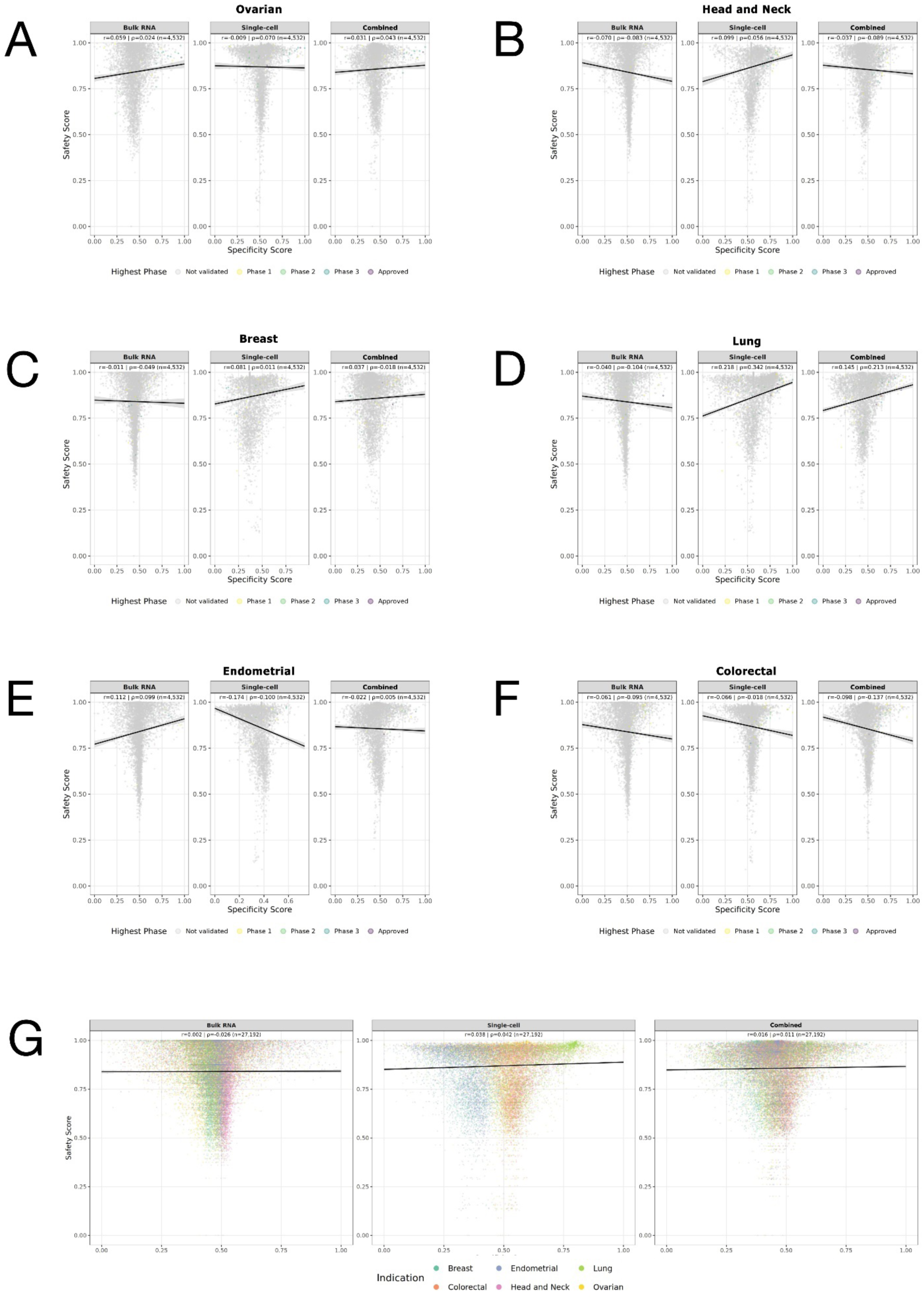
Relationship between tumor Specificity and normal-expression Safety scores across indications and expression-data modalities. Scatter plots show the relationship between Specificity and Safety scores across analyzed surfaceome genes. Scores were derived from bulk RNA-seq, single-cell RNA-seq, or combined bulk-and-single-cell evidence, as indicated above each panel. Black lines indicate linear regression fits. Pearson correlation coefficients (r) and Spearman rank correlation coefficients (ρ) are reported above each plot; correlation coefficients are presented as descriptive measures of association without hypothesis-testing p-values. **(A–F)** Indication-specific relationships for ovarian, head and neck, breast, lung, endometrial, and colorectal cancer, respectively, with 4,532 surfaceome genes evaluated per indication. Points represent individual candidate genes and are colored according to the highest clinical-stage threshold reached by the corresponding target; genes without clinical-stage validation are shown in gray. **(G)** Pooled analysis across all six indications, with points colored by cancer indication and each gene–indication pair treated as a separate observation (n = 27,192). Overall, Specificity and Safety showed weak linear and rank correlations, indicating that the two features capture largely distinct aspects of target expression.

